# The hepatic mitochondrial landscape

**DOI:** 10.64898/2026.07.03.736316

**Authors:** Jernej Vajda, Laura Cinc Curic, Uros Maver, Tomaz Martini

## Abstract

Mammalian energy homeostasis depends on coordinated metabolism across tissues, with the liver acting as a central hub for systemic energy balance and biosynthetic precursor supply. Hepatic mitochondria are integral to these functions, yet mitochondrial gene expression across liver microanatomical space remains incompletely defined. Here, we mapped the expression of genes supporting mitochondrial function at single-cell level across the feeding-fasting cycle and spatial gradients within the liver lobule in female and male mice. Hepatocyte subtypes exhibited pronounced mitochondrial divergence, including a non-canonical population characterised by low mitochondrial gene expression and reduced secretory protein production. We showed that higher periportal oxidative phosphorylation relies on an exceptionally high periportal mitochondrial transcript fraction that mirrors the mitochondrial-to-cell volume ratio, directly linking scRNA-seq read counts to mitochondrial content while also highlighting the limitations of the mitochondrial read fraction as a standalone scRNA-seq quality metric. Nuclear- and mitochondrial-encoded oxidative phosphorylation genes exhibited distinct zonation, revealing a lack of stoichiometric coordination at the transcript level. Ultrastructural reconstruction revealed zonation of mitochondrial architecture beyond size, including increased branching of periportal mitochondria. In humans, we recapitulated the higher periportal mitochondrial transcript abundance using spatial transcriptomics data. Mitochondrial gene expression rhythms, previously reported in mice, were also conserved in humans and peaked around the first meal in both species. Together, these findings reveal heterogeneity in hepatic mitochondrial biology and provide a framework for interpreting mitochondrial variation in liver single-cell studies.

## Introduction

Mammalian physiology relies on tightly coordinated energy metabolism across tissues. The liver plays a central role in systemic metabolic regulation, while also providing essential biosynthetic products to other organs and detoxifying harmful metabolites. It serves as a key energy buffer, particularly during fasting, feeding, and circadian fluctuations. These diverse and dynamic functions demand a highly adaptable and efficient mitochondrial network. Hepatic mitochondria, therefore, are central hubs for energy production, redox balance, and metabolic signaling.

The liver is especially well suited to govern the organism’s adaptation to times of fasting and feeding, as it not only senses recurring external metabolic cues but also has robust intrinsic circadian clock machinery that functions as a self-sustained molecular oscillator, allowing for the anticipation of recurring events, such as nutrient intake. Such anticipatory regulation is essential because relying solely on nutrient sensing would be inefficient given the inherent lag between signal detection and protein-mediated responses^1,2^.

Additionally, the liver is composed of repetitive functional units termed lobules. The outer layers of a lobule receive oxygenated and nutrient-rich blood via the hepatic artery and portal vein, respectively. The blood then flows towards the inner layers where deoxygenated blood is collected and extracted from within the lobule via the central vein. This organisation, further enhanced by signaling gradients originating from hepatic non-parenchymal cells, results in markedly different biochemical programmes in pericentral (PC) vs. periportal (PP) hepatocytes, a characteristic referred to as metabolic zonation ^1^.

Together, the metabolic versatility of the liver is shaped by its ability to perform specialised tasks depending on time of day, and by the ability to compartmentalise metabolism within hepatic microanatomy. Within this temporal and spatial framework, mitochondrial function is governed by both mitochondrial-encoded genes and a large repertoire of nuclear-encoded genes that coordinate mitochondrial biogenesis, dynamics, and metabolic output ^3^. While the biological roles of these gene sets are increasingly understood, less is known about how their expression varies at the single-cell level, within the architecture of the hepatic lobule, but also between sexes, factors known to influence liver metabolism.

Owing to the physiological role of mitochondria, hepatic mitochondrial dysfunction is implicated in a wide spectrum of diseases, including metabolic dysfunction-associated steatotic liver disease (MASLD), type 2 diabetes, and cardiovascular disease, as well as rare monogenic conditions that manifest early in life due to inherited defects in mitochondrial genes ^1,4–6^. With the emergence of gene therapy, numerous hepatic diseases could be corrected or managed. The liver is an optimal target for genetic manipulation, as intravenous delivery of adeno-associated viruses, which are commonly used vectors, primarily transduces the hepatic parenchyma ^7–9^. Emerging therapeutic approaches using chemically modified RNA also efficiently target and modulate liver function ^10^. The success of these therapies, however, requires a precise understanding of baseline hepatic gene-expression patterns. This knowledge is equally critical given the increasing availability of large-scale single-cell and single-nucleus RNA sequencing and proteomic datasets, which demand careful attention to fundamental liver biology. Owing to the high bioenergetic demands of hepatocytes, specific hepatocyte subtypes may be disproportionately and inadvertently removed during single-cell quality-control procedures, as elevated mitochondrial RNA content is commonly used as a proxy for poor cell quality or compromised membrane integrity.

The convergence of clinical need and technological capability now provides a timely opportunity to clarify the spatiotemporal and sex-dimorphic regulation of hepatic mitochondrial function. Here, we integrated deeply sequenced and spatially reconstructed scRNA-seq in female and male mouse liver samples ^11^ with human spatial transcriptomics data ^12^ and scanning electron microscopy to examine mitochondrial biology along the lobular axis, and revisited mitochondrial rhythmicity in bulk RNA-seq datasets. These complementary approaches revealed several previously unresolved aspects of hepatic mitochondrial biology, including increased branching of PP mitochondria, spatially organised stoichiometric imbalances among oxidative phosphorylation genes, candidate regulators of zonated mitochondrial function, conserved zonation of mitochondrial transcript abundance in mouse and human liver, with a marked PP increase in mitochondrial transcripts in both species, and a relationship between mitochondrial read fractions in scRNA-seq and cellular mitochondrial content. Together, these insights provide a framework for interpreting large-scale hepatic datasets and mitochondrial dysfunction in disease.

## Results

### Functional hepatocyte subtypes are defined by mitochondrial divergence

Hepatocytes exhibit substantial variation in mitochondrial transcript abundance beyond their established zonal and temporal transcriptional heterogeneity ^11,13,14^. We previously identified a subset of cells within the principal hepatocyte cluster of our liver scRNA-seq dataset with a different overall transcriptomic signature, including a markedly attenuated expression of mitochondrially encoded cytochrome c oxidase subunit I (*mt-Co1*), a core component of the mitochondrial respiratory complex IV. Several observations support the potential biological relevance of this population. Clonally derived cytochrome c oxidase-deficient hepatocyte patches have been identified in the human liver, extending from portal tracts towards central veins, consistent with clonal expansion from long-lived PP cells ^15^. Moreover, somatic mutations in *mt-Co1* exhibited signatures of selection during ageing, while experimental complex IV deficiency induced hepatic steatosis and stimulated transdifferentiation of biliary epithelial cells during hepatic regeneration ^16,17^. Therefore, we sought to characterise *mt-Co1*⁻ entities at single-cell transcriptomic resolution. We first refined cell assignment to the *mt-Co1*⁻ cluster based on a two-component Gaussian mixture model (Fig. 1A).

**Figure 1:**
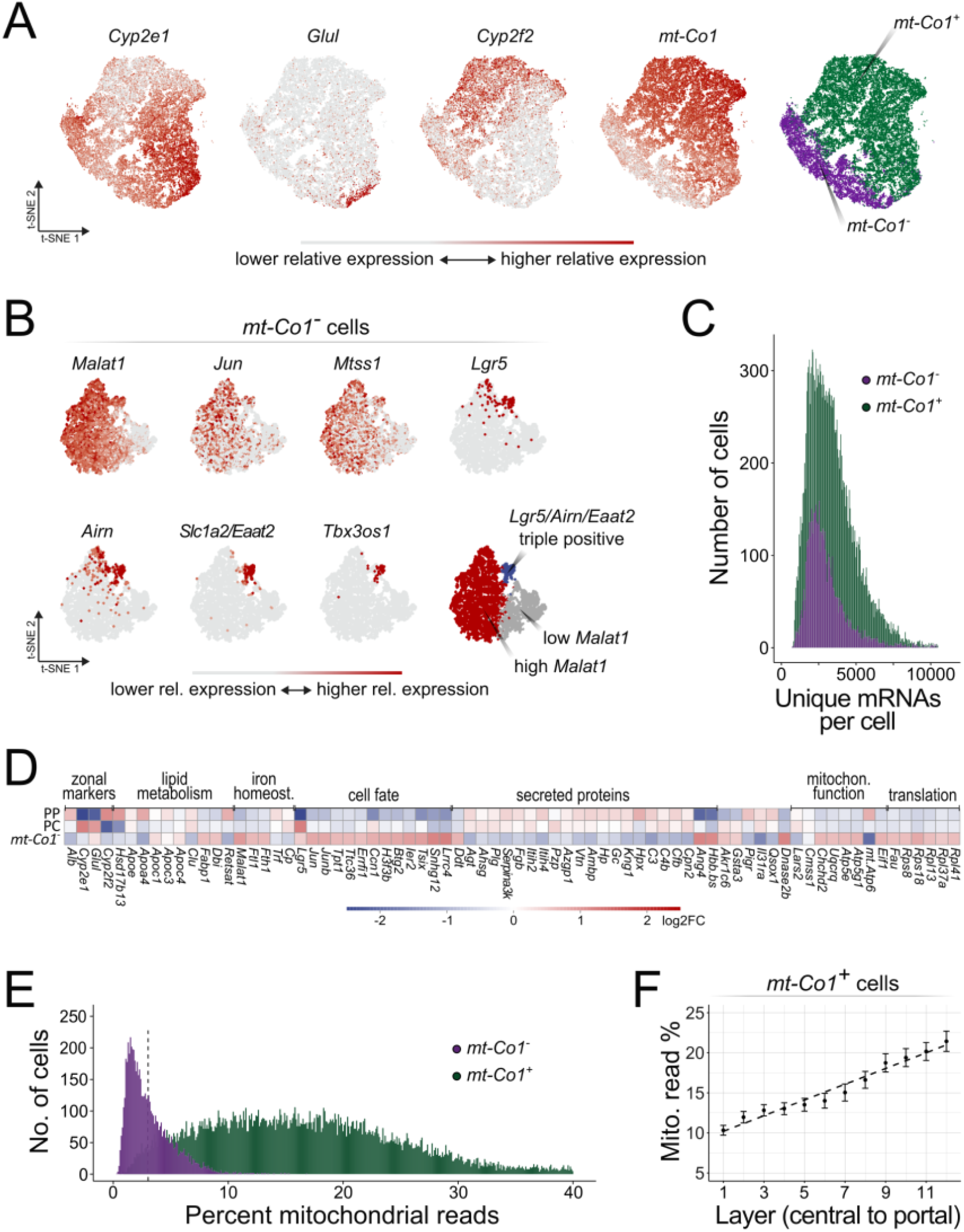
Gross division of hepatocyte identity aligns with differences in mitochondrial read percent in 129S mice. (A) Dimensionality reduction of hepatic scRNA-seq data. For contextualisation of hepatocyte zonation, the left three panels are shown for spatial reference and were adapted from ref. ^11^. After removal of nonparenchymal-cell clusters from the liver scRNA-seq data, cells in the hepatocyte cluster split by zonal identity (seen by expression gradients of the zonation markers Cyp2e1, Glul and Cyp2f2). Cells split by mt-Co1 expression in the axis perpendicular to the one corresponding to zonation. The variability in mt-Co1 expression was utilised to split the cells into mt-Co1^+^ and mt-Co1^⁻^ (right-most panel; n = 12 mice). (B) Reclustering of mt-Co1^⁻^ cells reveals three further subclusters, one of which is characterised by Lgr, Slc1a2, Airn and Tbx3os1 expression. (C) As a quality control, the distribution of unique RNA molecules (unique molecular identifiers or UMIs) was assessed for both mt-Co1^+^ hepatocytes and mt-Co1^⁻^ cells, showing comparative distribution profiles. (D) Heatmap showing gene expression in PC and PP hepatocytes and mt-Co1^⁻^ cells compared to mean hepatocyte mRNA levels of each individual gene, with 2 most PC and PP virtual layers of hepatocytes analysed, respectively. mt-Co1^⁻^ cells show transcriptomic specifics, including reduced expression of genes encoding blood-borne proteins. (E) The mt-Co1^+^ hepatocytes and mt-Co1^⁻^ cells show a different total mitochondrial read percent, with a wide distribution of reads in canonical hepatocytes. (F) Aggregated visualisation of mitochondrial read fraction across the porto–central axis, based on data from ref. ^11^. In the canonical mt-Co1^+^ hepatocytes, the percent of mitochondrial reads shows an increase in the pericentral to periportal direction.

Reclustering of the *mt-Co1*⁻ cells split these into three further groups, two of which exhibited higher *Malat1*, *Mtss1* and *Jun* expression. One of these two subclusters was *Lgr5*-, *Airn*-, *Slc1a2*- and *Tbx3os1*-positive (Fig. 1B). *Lgr5* is a well-known stem cell marker associated with regenerative responses in the liver ^18^, and hepatic *Lgr5*⁺ cells can give rise to hepatocyte organoids and are activated during liver damage ^19,20^. Notably, *mt-Co1*⁻ and *mt-Co1*^+^ groups showed a similar distribution of detected unique mRNAs per cell (Fig. 1C) and intronic reads (SFig. 1A), arguing against cell lysis or substantial RNA loss as the explanation for the *mt-Co1*⁻ state. Together with the tissue evidence described above, these findings support the existence of a genuine *mt-Co1*⁻ population rather than a technical artifact.

Differential expression analysis between *mt-Co1*⁻ and *mt-Co1*^+^ clusters revealed a reduced expression of multiple mitochondrial-encoded genes and mature hepatocyte markers, including *Alb*, in *mt-Co1*⁻ cells (Fig. 1D, SFig. 1C, S. Table 1-3). The reduced expression of mitochondrial DNA-encoded transcripts contrasted with the expression of nuclear-encoded genes supporting mitochondrial function, many of which exhibited unaltered or even modestly elevated expression in the *mt-Co1*⁻ cluster. *mt-Co1*⁻ cells also showed attenuated expression of other liver-derived blood-borne products beyond *Alb*, such as *Apoa4* and components of the complement and coagulation pathways, and showed no consistent PC or PP signatures across canonical zonal markers. Conversely, *mt-Co1*⁻ cells exhibited an upregulated expression of immediate early genes (*Jun*, *JunB*, *Ier2*, *Btg2*) and chromatin- or splicing-related regulators (*H3f3b*, and the lncRNAs *Malat1* and *Snhg12*). As relative expression is affected by the depletion of abundant hepatocyte transcripts, the higher expression of these gene groups should be interpreted with caution. Functional studies will be required to further examine the physiological importance of *mt-Co1*⁻ entities.

Splitting the cells by *mt-Co1* expression also revealed the variability of total mitochondrial reads in both *mt-Co1*⁻ cells and *mt-Co1*^+^ canonical hepatocytes (Fig. 1E), with a particularly wide distribution of mitochondrial reads in *mt-Co1*^+^ hepatocytes, consistent with our previous report ^11^.

### Liver zonation is mirrored by mitochondrial transcript abundance and morphological changes

To further explore biological variability in canonical hepatocytes, we performed spatial reconstruction by digitally repositioning individual hepatocytes along the PC–PP axis ^11^, and focused specifically on mitochondrial features of canonical hepatocytes and the comparison of PC and PP hepatocytes with *mt-Co1*⁻ cells (Fig. 1D). Given our emphasis on mitochondrial biology, which can be influenced by strain-specific genetics, experiments were conducted in 129S6/SvEv (henceforth 129S) rather than commonly used substrains of C57BL/6 mice, as these carry an *Nnt* loss-of-function mutation that impairs mitochondrial redox balance and increases oxidative stress, thereby reducing translational relevance to human metabolism ^21–23^.

Despite the well recognised differences between PC and PP canonical hepatocytes ^14,24^, these populations were qualitatively more similar to each other than to the *mt-Co1*⁻ entities. However, canonical hepatocytes exhibited a transcriptomic gradient of mitochondrial gene expression along the PC–PP axis, as previously observed in our dataset ^11^. A quantitative analysis revealed that mitochondrial transcripts comprised about 10% of the pericentral hepatocyte transcriptome, increasing to a mean of 20% periportally (Fig. 1F, SFig. 1D). To assess the contribution of mitochondrial abundance and organisation to this transcriptional gradient, we examined mitochondrial ultrastructure.

The increase in mitochondrial reads toward PP areas correlated with distinct mitochondrial morphological changes. Scanning electron microscopy (SEM) performed after the feeding phase in mice revealed larger mitochondria that occupied a greater area in PP compared to PC hepatocytes in individual planes (Fig. 2A), and mitochondrial reconstructions in 3D revealed larger mitochondrial volumes in PP hepatocytes (Fig. 2B-C). Although earlier estimates suggested that PP mitochondria may be larger ^25,26^, we validated this difference across larger tissue areas and biological replicates, in both female and male mice (Fig. 2C-D, SFig. 2). Additionally, we also observed that PP mitochondria exhibited more heterogeneous morphology than PC mitochondria, revealing an additional dimension of mitochondrial zonation beyond average size (Fig. 2B). Multiple PP mitochondria appeared branched in our 3D reconstructions rather than spherical or cylindrical. Accordingly, distributions of volume as a function of mitochondrial sphericity differed markedly between PC and PP regions (Fig. 2C). PC hepatocytes occupied a tight, low-volume morphospace, while PP hepatocytes spanned a wider morphometric range with a distinct high-volume, low-sphericity tail, suggesting that more voluminous mitochondria are those exhibiting a higher degree of branching, possibly due to increased dynamic remodelling. This observation contrasts with the earlier description of predominantly rounded, short PP mitochondria ^25^, and may inform models of zonal mitochondrial dynamics. The coexistence of distinct mitochondrial architectural states within PP hepatocytes raises the possibility of differences in mitochondrial biogenesis, turnover or function.

**Figure 2:**
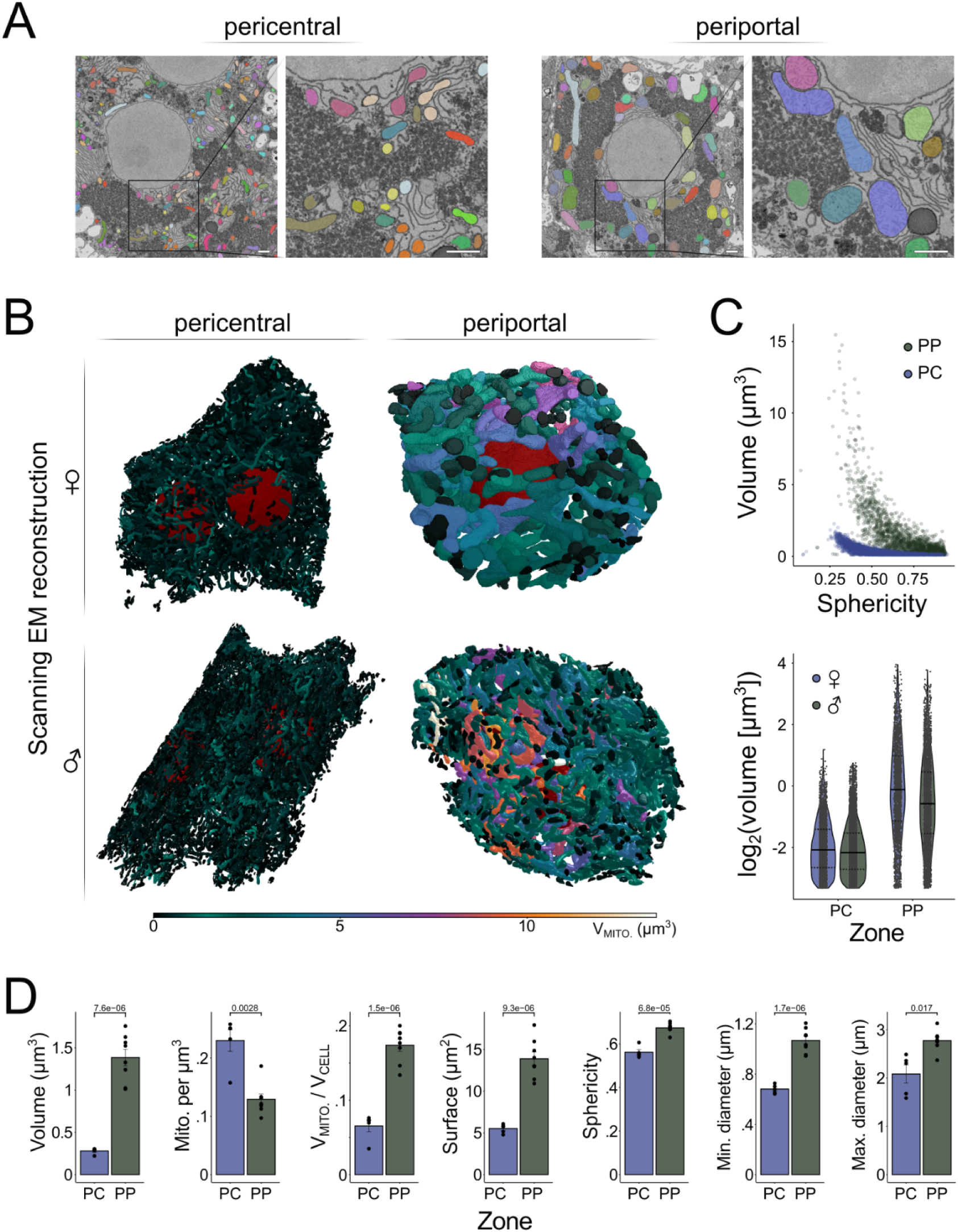
Morphological characteristics of pericentral and periportal hepatocytes in 129S mice. (A) Mitochondrial segmentation examples of PC and PP hepatocytes, an individual 2D plane of SEM is shown. In the PC hepatocyte, smaller mitochondria are visible, with discrete dark granular areas corresponding to glycogen. In the PP hepatocyte, mitochondria appear thicker and occupy larger surface areas, and large glycogen stores are visible throughout the entire cell. Scale bars represent 1 μm. (B) Spatial reconstructions of mitochondria from SEM of PC and PP hepatocytes of female and male mice, representative cells are visualised. PP mitochondria appear thicker and occupy a greater relative volume of the cell. Due to differences in PC and PP cell sizes, the individual cells are not represented at the same scale. (C-D) Quantitative assessment of the mitochondrial 3D reconstructions. (C) Quantifications of individual mitochondria, each dot represents one mitochondrion. Top panel, relationship between the absolute mitochondrial volume and sphericity, for PP and PC mitochondria. In the volume vs. sphericity space, PP and PC mitochondria separate into clusters. Bottom panel, absolute per-mitochondrion volume distributions in PC and PP hepatocytes from female and male mice. In the violin plots, solid lines depict the median, and dotted lines the lower and upper quartile. Both PC and PP hepatocytes show wide mitochondrial volume distributions, and PP mitochondria are generally larger (n = 14894 mitochondria; 8 samples; 4 mice). (D) Quantification of cell means, each dot represents the mean value of an individual cell, bars indicate mean values across cells. PP mitochondria have larger volumes, increased thickness (min. Feret diameters) and length (max. Feret diameters), and higher sphericity compared to PC mitochondria. Sphericity is calculated as the ratio of the surface area of a sphere with the same volume as the object to the surface area of the object. Error bars represent SE, p-values are shown (Student’s t-test). The tissue was analysed at ZT 2 (2 h after lights were switched on in the housing facility), in the fed state (n = 8 regions; 4 mice).

Extending these observations across individual hepatocytes and biological replicates, in female and male mice, quantitative 3D morphometry revealed zonated differences across multiple features of mitochondrial architecture beyond mitochondrial volume (Fig. 2D, SFig. 2). PP mitochondria had greater minimum Feret diameters, reflecting increased mitochondrial thickness. On average, the sphericity (a metric of overall shape compactness) of PP mitochondria was higher compared to PC mitochondria, likely due to the trend of greater minimum Feret diameters. PP mitochondria also exhibited greater lengths, represented as higher maximum Feret diameters.

Moreover, our reconstruction of larger tissue volumes allowed us to compare mitochondrial number and volume density between regions. PC hepatocytes showed a higher number of individual mitochondria per cell volume. Yet, the relative volume of mitochondria per cell volume was markedly higher periportally (Fig. 2D), with slightly less than 10% of the cell volume occupied by mitochondria pericentrally and just below 20% periportally. Remarkably, these values closely mirrored the zonated mitochondrial read fraction (Fig. 1E), despite being derived from an independent modality, suggesting that mitochondrial transcript abundance may provide a quantitative proxy for mitochondrial content in hepatocytes. Notably, mitochondrial reads are stable throughout the day (SFig. 1D) ^27^. Since PP hepatocytes are known for their production of blood-borne proteins, a secretory role that entails high metabolic demand, the mitochondrial enrichment in PP hepatocytes may sustain this specialised biological role of governing the circulatory proteome ^1^.

### Spatiotemporal compartmentalisation of catabolism and anabolism in female and male mice

The mammalian liver compartmentalises metabolic processes across both time (Fig. 3A) and microanatomical space. Pioneering spatially reconstructed scRNA-seq studies in male C57BL/6J mice captured mitochondrial genes as individual examples or within pathway annotations ^14,24^. A mitochondria-focused reference atlas resolving feeding-fasting state, lobular position and, crucially, sex at single-cell resolution is therefore lacking. All three variables are known to affect hepatic physiology, and without such an atlas, mitochondrial read fractions and gene programmes in hepatic single-cell datasets are difficult to interpret.

**Figure 3:**
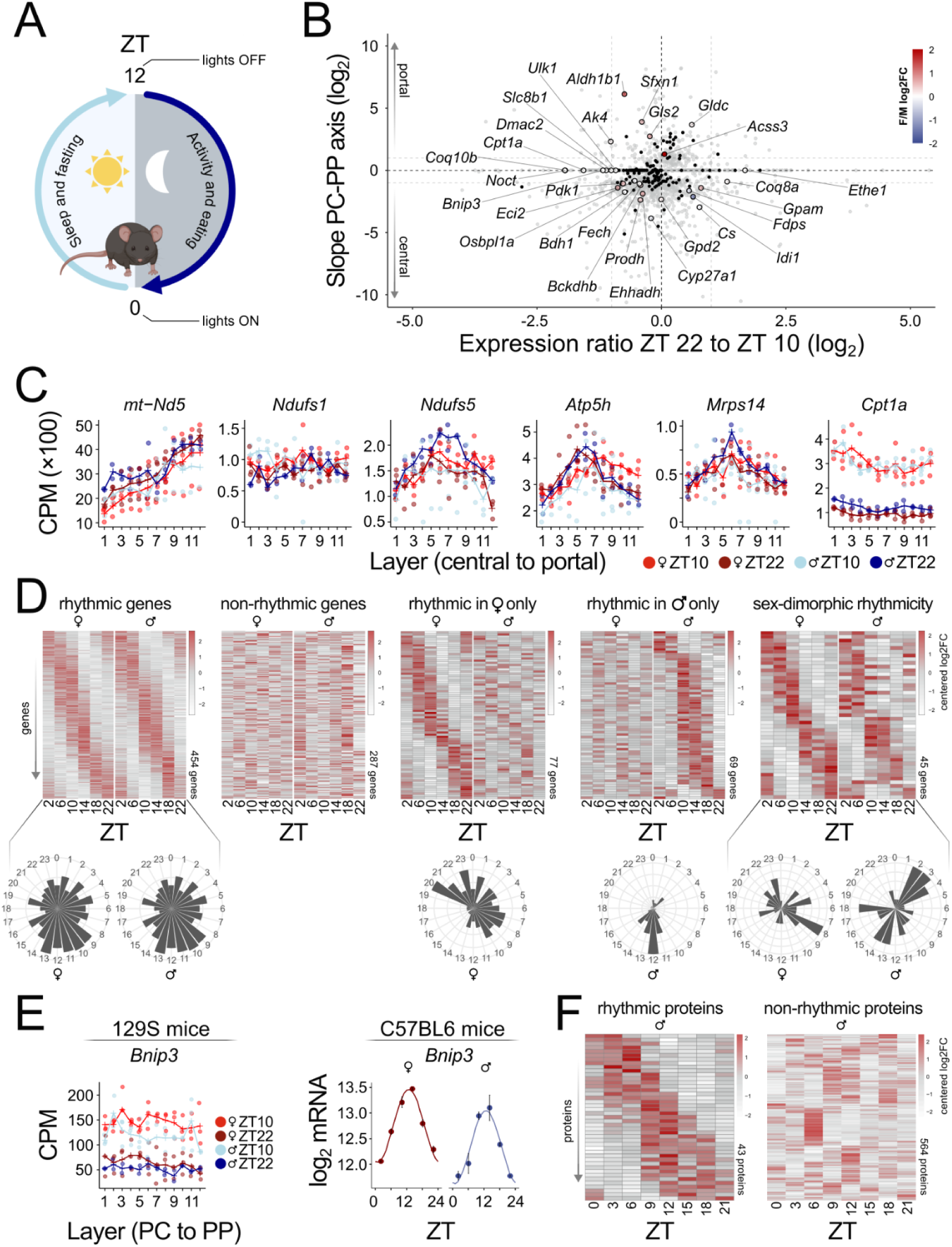
Spatiotemporal compartmentalisation of mitochondrial-driven catabolism and anabolism. (A) Schematic representation of the murine light-dark cycle and associated behavioural states. Lights turn on at ZT 0 and off at ZT 12, dividing the 24-h cycle into a light phase (sleep and fasting) and a dark phase (activity and eating). (B–C) Spatial reconstruction of hepatocytes allows the plotting of gene expression in function of virtual hepatocyte layers in the PC-PP direction, with 1 corresponding to the most PC layer. The experiments were conducted in 129S mice (n = 11). (B) A summary of the slope of the spatial expression plots (negative slopes characterise central profiles) as a function of the expression ratio between ZT 22 and ZT 10. Gray dots represent all detected genes, mitochondrial-function associated genes are shown in black. A selected subset of mitochondrial genes with pronounced physiological and pathophysiological importance includes explicit gene names and emphasised points, which are coloured according to the female to male expression ratio. The ratios are plotted on the log_2_-scale. Genes in the corners are highly zonated and strongly regulated by time. (C) Reconstructed spatial profiles of gene expression in 12 virtual hepatocytes layers, 1 corresponding to the most PC layer. Counts are normalised to the total counts of cytoplasmic unique molecular identifiers (UMIs; uniquely detected mRNAs) and represented as counts per million (CPM; n = 11; n = 2–3, per condition). Examples include mitochondrial and nuclear-encoded genes, which are differently affected by zone and time, including genes with midlobular peaks, such as Ndufs5 and Atp5h. Genes that peak midlobularly have small or non-significant PC-PP slopes and are elusive in slope-based summary plots. (D) Bulk RNA-seq, conducted in C57BL/6J mice, performed at multiple time-points around the clock, allowed the circadian analysis of mitochondrial-associated genes (MitoCarta 3.0). Heat maps of normalised rhythmic mRNA levels in the liver are shown, with each row corresponding to a gene, with per-gene mean-centered expression. Rhythmic shifts from daily mean expression levels of a particular gene are depicted as log_2_-fold changes (log2FC). Expression amplitudes were capped at –3 and 3, respectively. Genes were classified as rhythmic in female and male mice or non-rhythmic in either sex, while some genes were differentially rhythmic in female and male mice, with rhythmicity either in only one sex or with a phase shift between females and males. Approximately half of mitochondrial function associated genes exhibit profound rhythmicity with a peak at the fasting-feeding transition in both sexes (left-most panel). The distribution of peak expression phases (acrophases) for the rhythmic genes demonstrated that mitochondrial function is strongly temporally programmed to follow fasting-feeding cycles in mice. (E) Spatial mRNA expression profiles in counts per million (CPM) reveal that Bnip3 is non-zonated, but its expression levels are affected by time-of-day in both female and male mice (129S mice; n = 11). Bulk mRNA expression confirms rhythms in Bnip3 in female and male livers (C57BL/6 mice; n = 24). (F) Proteomics rhythmicity assessment from ref. ^2^ shows that in male mice roughly a tenth of mitochondrial proteins show robust rhythmicity.

The spatial reconstruction of our scRNA-seq dataset from female and male 129S mice, sampled at two physiologically distinct time-points, revealed how genes associated with mitochondrial function vary across the lobule, either on a per-gene basis (Fig. 3B, SFig. 3A-B) or when grouped by mitochondrial subcompartment (SFig. 3C). The resulting visualisations provide a resource for the zonation, temporal regulation and sex-associated expression of all mitochondrial genes responsive to at least one of these three variables (*i.e*., a log_2_-fold change of > 1 in at least one dimension; SFig. 3A-B). Overall, the data suggest that mitochondrial gene expression is spatially zonated, but also affected by time of day, while a subset of genes exhibits sex-biased expression (Fig. 3B). A key pattern emerging from this analysis was a consistent divergence between mitochondrial- and nuclear-encoded components of the oxidative phosphorylation (OXPHOS) system, consisting of electron transport chain (ETC) genes and the ATP synthase complex (SFig. 4). Mitochondrially encoded subunits are enriched periportally, whereas their nuclear-encoded counterparts tend to be non-zonated or peak midlobularly. This disconnect was observed across all OXPHOS complexes with mitochondrially encoded subunits and suggests that spatial regulation of oxidative phosphorylation capacity is primarily driven by the periportally enriched mitochondrial genome, corresponding to the higher partial oxygen pressure in PP hepatocytes.

**Figure 4:**
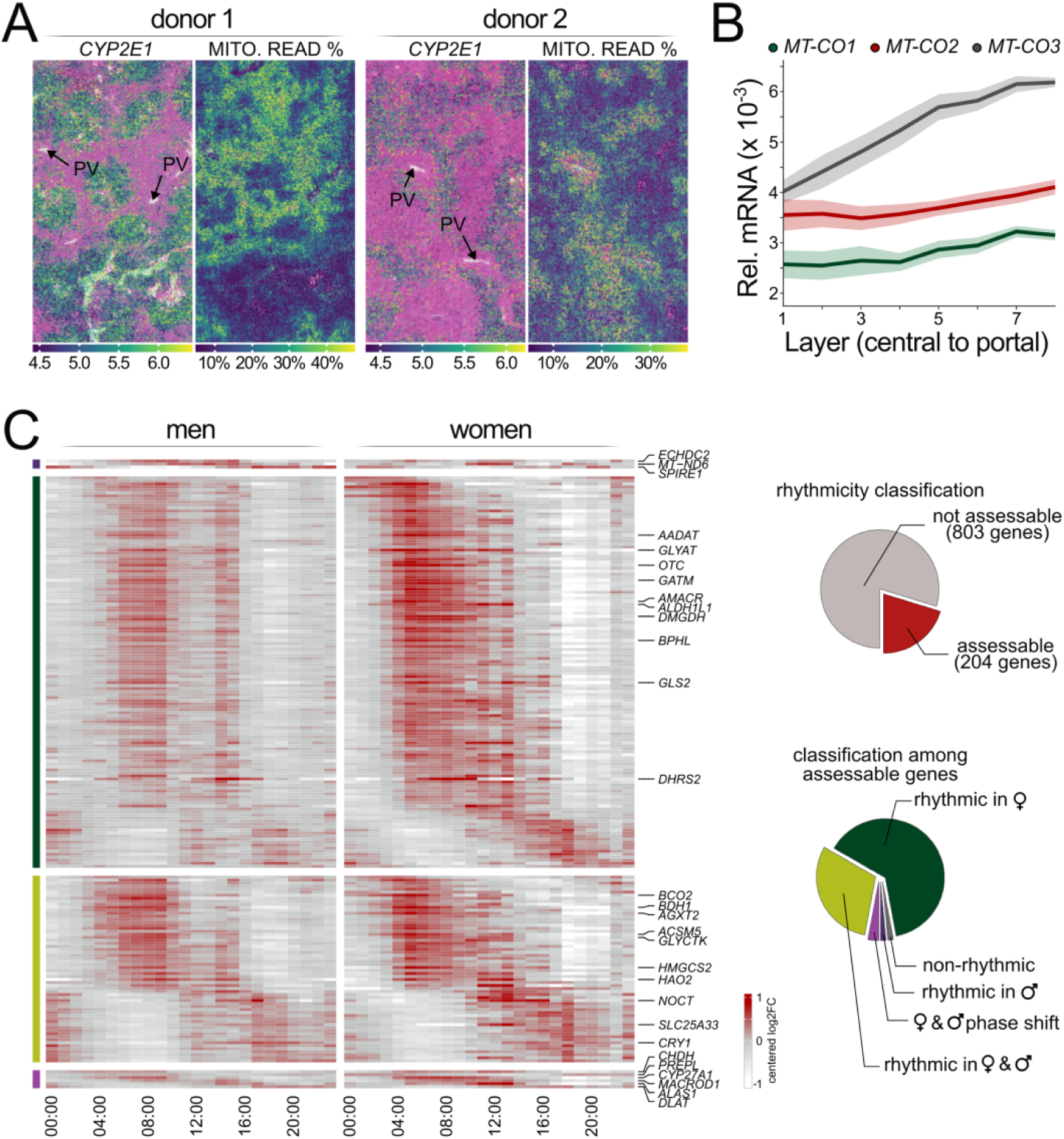
Human liver mitochondrial transcripts are spatially zonated and sex-dependently rhythmic. (A) Spatial transcriptomics, reanalysed after ref. ^12^, allows the visualisation of gene expression on hematoxylin-eosin-stained tissue slices. CYP2E1 (left panel) is shown as a representative PC zonation marker. Mitochondrial transcript abundance is antizonated with CYP2E1 expression, suggesting a PC-PP mitochondrial read percent gradient. Colour scales indicate log-transformed CYP2E1 expression and mitochondrial read fraction per spatial bin. (B) Relative mRNA abundance of MT-CO1, MT-CO2 and MT-CO3 increases from central to portal layers, reanalysed after ref. ^12^. Lines indicate mean relative mRNA abundance and shaded regions indicate standard errors of the mean. (C) Analysis of rhythmic mRNA expression for mitochondrial genes in human bulk RNA-seq data from the GTEx project. Heatmaps show genes assessed as rhythmic in men only, women only, rhythmic in both men and women, or rhythmic in both men and women with a phase shift along the time axis. For each model, the 10 genes with the highest amplitudes are explicitly annotated, unless the model has less than 10 genes. Pie charts show the proportion of mitochondrial genes assessable for rhythmicity and the distribution of rhythmicity classes among assessable genes (n = 224; 160 men, 64 women).

Consistent with this observation, mitochondrial complex I (NADH dehydrogenase) genes displayed a particularly strong PP increase, while nuclear encoded complex I (*Nduf*) genes exhibited non-zonated or midlobularly increased mRNA expression (Fig. 3C, SFig. 3A-B, SFig. 4). Complex II (*Sdh*) genes, entirely nuclear-encoded, did not exhibit a periportal bias (SFig. 4). These patterns suggest PP hepatocytes specialise in NADH-driven oxidative phosphorylation and ATP generation, whereas succinate oxidation via complex II serves a uniform, possibly housekeeping role. Nuclear complex III genes are largely non-zonated or mildly midlobularly enhanced in expression, except for the periportally biased mitochondrial *mt-Cytb* (SFig. 3A-B, SFig. 4). Complex IV genes *mt-Co1*–*3* were strongly periportal, while nuclear *Cox* genes again peaked midlobularly (SFig. 3A-B, SFig. 4). Complex V genes *mt-Atp8* and *mt-Atp6* were periportally biased, contrasting with non-zonated or higher midlobular levels of specific *Atp5* nuclear genes, notably *Atp5h* (Fig. 3C, SFig. 3A-B, SFig. 4). Together, the results suggest that zonated changes in OXPHOS gene expression do not follow a stoichiometric balance, although this imbalance is inferred at the transcript level only and does not describe potential differences in translation or protein stability.

While the ETC represents a core mitochondrial function, metabolic regulation often occurs at the pyruvate branch point, where multiple pathways intersect. Pyruvate can enter the tricarboxylic acid (TCA) cycle to generate reduced flavin adenine dinucleotide (FADH_2_) and reduced nicotinamide adenine dinucleotide (NADH), or be diverted to gluconeogenesis, de novo fatty acid synthesis, or ketone body production. *Dlat* encodes the catalytic subunit of the pyruvate dehydrogenase complex, which converts pyruvate to acetyl-CoA, and exhibits a strong pericentral enrichment. *Pdk1*, which phosphorylates and inhibits this complex, increases at the fasting–feeding transition, particularly in pericentral hepatocytes and females, suggesting spatiotemporal modulation of pyruvate flux into downstream metabolism (Fig. 3B, SFig. 3A-B). Moreover, *Hmgcs1*, the cytosolic 3-hydroxy-3-methylglutaryl-CoA synthase, rose toward the end of feeding and became pericentrally enriched, whereas it was non-zonated at the end of fasting, indicating a shift from ketogenic to mevalonate (and hence cholesterol) synthesis after feeding. Conversely, *Bdh1*, encoding the mitochondrial enzyme interconverting acetoacetate and β-hydroxybutyrate, exhibited modest pericentral enrichment and rose at the fasting–feeding transition, consistent with the role of β-hydroxybutyrate in systemic food anticipation ^9^. Within the TCA cycle, *Cs* (citrate synthase) was pericentrally expressed and more highly expressed at the feeding–fasting transition (Fig. 3B, SFig. 3A-B). Notably, multiple key transporters and enzymes involved in mitochondrial metabolism exhibited sex-dimorphic expression, including genes involved in lipid homeostasis (SFig. 3A-B).

We next sought transcriptional correlates of the zonated differences in mitochondrial ultrastructure. Genes involved in mitochondrial fusion, fission and mitophagy showed no clearly identifiable expression pattern that explained the observed morphological differences. These processes may instead be regulated post-transcriptionally. However, our results revealed markedly elevated PP expression of *Ak4* (Fig. 3B), a potential contributor to phenotypic differences between PC and PP mitochondrial structure. *Ak4* has been implicated as a driver of mtDNA synthesis ^28^, while *Ak4* knockdown in mice resulted in higher numbers of mitochondria ^29^. Indeed, we observed a larger number of smaller mitochondria pericentrally, where the expression of *Ak4* was markedly lower. Yet, functional experiments involving *Ak4* were performed in distinct biological contexts, and its contribution to mitochondrial ultrastructure in hepatocytes remains unknown.

Although our scRNA-seq analysis resolved spatiotemporal expression, it could not capture complete daily gene expression profiles. Bulk RNA-seq (or similar approaches) time courses sample the full day densely, but have mostly examined mitochondrial genes as part of genome-wide analyses ^30,31^. We therefore performed a targeted reanalysis of mitochondrial gene rhythmicity in female and male mice ^27^, complementing our spatial data. This analysis primarily served as a mouse benchmark for our subsequent comparison with human liver rhythms. However, the systematic analysis also revealed that roughly half of the analysed mitochondrial genes exhibited rhythmic expression in both sexes, while additional genes were rhythmic only in females or males, or displayed sex-specific phase shifts, broadly consistent with earlier liver chronotranscriptomic work (Fig. 3D) ^27^. Most genes, with equivalent expression rhythms in both sexes, peaked around the fasting-to-feeding transition, possibly in anticipation of nutrient influx. Among these were also regulators of mitophagy, including *Bnip3*, whose rhythmic expression was observed across two mouse strains (Fig. 3E), consistent with previous observations of its temporal regulation and in line with the proposed temporal alignment of mitophagy related pathways with the anticipated increase in mitochondrial workload upon feeding ^32^. Enhanced metabolic flux may increase mitochondrial stress and the need for tight control of mitochondrial turnover.

In our mitochondrial focused reanalysis of hepatic proteomics data ^2^, the rhythmicity observed at the mRNA level was only partially reflected at the level of the proteome, with robust rhythmicity detected for fewer than a tenth of proteins. This indicates a substantial uncoupling between rhythmic mitochondrial transcripts and steady-state protein abundance. However, limited rhythmicity in protein abundance does not necessarily indicate limited rhythmicity in mitochondrial function, as enzyme activity may also vary through rhythmic post-translational modifications and metabolite availability ^33^.

### Mitochondrial landscape in human livers

Differences between mice and humans in hepatic mitochondrial function may present a challenge when translating research findings from animal models to human biology. To evaluate whether the mitochondrial zonation observed in mouse liver is conserved in humans, we re-analysed an existing human spatial transcriptomics dataset from healthy liver donors ^12^, showing a profound effect of zonation on mitochondrial read percentage, with markedly higher mitochondrial reads in PP compared to PC hepatocytes (Fig. 4A; SFig. 5A). This pattern closely recapitulated the mitochondrial gradient observed in mice, establishing PP enrichment of mitochondrial transcripts as a conserved feature of mammalian liver zonation. The PP increase in total mitochondrial reads was also reflected in zonated *MT-CO1/2/3* expression profiles ^12,34^ (Fig. 4B).

Moreover, similarly to the results from mice, analysis of human bulk liver transcriptomes from the Genotype-Tissue Expression (GTEx) project ^35,36^, using circadian phase inference ^37^, revealed rhythmicity of mitochondrial- and nuclear-encoded genes involved in mitochondrial function. Despite the inherently higher biological variability in human data compared to mice, approximately a fifth of the mitochondrial genes, for which rhythmicity could be assessed, showed robust rhythmic gene expression (Fig. 4C). Most rhythmic genes tend to peak around the first meal of the day, equivalently to what we observed in mice. Moreover, more than half of the genes that could not be statistically classified for rhythmicity displayed rhythmic-like temporal profiles, suggesting that statistical classification may underestimate the extent of mitochondrial rhythmicity within the biologically variable human cohort (Fig. 4C, SFig. 5B). A number of genes, which exhibited rhythmic-like profiles but did not meet the statistical criteria for rhythmicity, peaked later, near the middle of the human active phase (SFig. 5B). Conversely, the remainder of the rhythmically non-assessable genes displayed non-rhythmic-like expression patterns, although many also showed a modest increase in expression around mid-day (SFig. 5B). This trend prevented robust classification of these genes as non-rhythmic and likely reflects a common time-of-day bias rather than bona fide rhythmic regulation.

Mitochondrial rhythmicity in the human liver was, as in mice, sex-dependent, with rhythmic genes separating into shared and sex-specific classes (Fig. 4C). However, contrary to the results from mice, human data suggested markedly higher rhythmicity of mitochondrial genes in female liver. For those genes, the male expression pattern showed only modest differences in expression throughout the duration of a single day.

Among the rhythmic transcripts with direct clinical relevance, *OTC* stood out as a particularly notable example, as it encodes ornithine transcarbamylase, an enzyme whose deficiency results in the most common urea cycle disorder in humans, a severe monogenic liver disease characterised by impaired hepatic ammonia detoxification and life-threatening hyperammonaemia. The strong rhythmicity of *OTC* mRNA in human liver, particularly in women, is noteworthy, as it shows that a clinically actionable metabolic disease gene is not expressed as a static hepatic transcript, but is embedded within the temporal program of liver physiology. This observation raises the possibility that constitutive episomal expression following gene addition therapy could have unanticipated long-term consequences if it disrupts the normal temporal regulation of hepatic nitrogen metabolism.

## Discussion

Recent technological advances have accelerated the development of methods for assessing cell function at the single-cell level, including in liver tissue. However, despite significant progress in understanding hepatic physiology at the spatiotemporal level, the mitochondrial landscape is often underappreciated, while being critical for proper data interpretation and quantitative normalisation in omics approaches, including scRNA-seq. An unbiased and comprehensive analysis of the hepatic mitochondrial landscape can therefore underpin future systematic analyses of hepatic tissue, or allow the reanalysis of previous studies.

Building on earlier efforts to optimise cell inclusion criteria through mitochondrial read percentage thresholds ^13,14^, our subsequent work ^11,38^ analysed mitochondrial read zonation across the liver lobule, revealing a progressive increase from PC to PP hepatocytes, while also providing initial scanning EM-based reconstructions of mitochondrial morphology differences in PC vs. PP hepatocytes. Later studies further characterised mitochondrial features under different physiological states, including approaches combining pseudobulk proteomics with EM ^25^ and analyses of animals sampled during the circadian fasting phase compared to prolonged food deprivation, the latter resulting in rounded mitochondrial morphology likely reflecting extensive mitophagy induced by severe nutrient loss ^26^. Here, we extended our initial transcriptomic analysis towards mitochondrial features, separating hepatocytes based on their mitochondrial divergence, either based on *mt-Co1* expression or zonal position. Moreover, with our 3D data from female and male mice, and experiments capturing larger volumes and biological replicates, we showed greater mitochondrial structural diversity as previously reported, including highly branched periportal mitochondria, especially those of higher volumes. Together with the validation of our observations in human liver, these datasets now provide a rich repository for prospective studies focusing on differential mRNA and protein abundance in mitochondrial-associated pathologies.

Notably, the divergence between mitochondrial- and nuclear-encoded gene expression along the zonation axis for OXPHOS complexes suggests that enriched PP oxidative capacity is likely not governed by nuclear transcriptional regulation, but is primarily driven by the enrichment of the mitochondrial genome. Accordingly, our data show that in mice mitochondrial RNA reads in hepatocytes correlate with the relative volume occupancy of mitochondria. This relationship may be relevant to mitochondrial pathologies, as inherited defects in mitochondrially encoded ETC subunits, which are enriched periportally, could result in zonal differences in the manifestation of mitochondrial defects. Conversely, nuclear-encoded complex deficiencies may manifest more uniformly across the lobule, a distinction that could inform histopathological interpretation in mitochondrial hepatopathies. However, ETC activity is unlikely to scale linearly with the abundance of every transcript. In multiprotein complexes and coupled reaction chains, functional output may depend disproportionately on limiting subunits, assembly efficiency, complex stability, substrate availability or downstream biochemical bottlenecks, whereas other components may only need to be maintained within a permissive functional range. These features may help contextualise why mitochondrial encoded transcripts show stronger zonation than many nuclear encoded ETC transcripts.

Our results from mice and humans also suggest that the mitochondrial landscape should not be studied as a static biological entity, but rather a temporally programmed and dynamic system, adding a temporal dimension to the understanding of mitochondrial dysfunction. Circadian misalignment associated with shift work, sleep disruption, or obesity-driven clock dysfunction may therefore impair hepatic mitochondrial homeostasis independently of nutrient availability. Conversely, the data raise the possibility that time-restricted feeding strategies, which reinforce circadian feeding-fasting cycles ^39^, could partly exert their hepatoprotective effects through the restoration of rhythmic mitochondrial function, including mitophagy.

In mice, we observed a substantial uncoupling between mRNA and protein rhythmicity. However, the limited rhythmicity in steady-state protein abundance does not imply that rhythmic mRNA expression is physiologically irrelevant. Rather, rhythmic transcripts may support time-of-day dependent protein synthesis, whereas protein abundance is further shaped by translational regulation, protein stability and degradation. Moreover, detectable protein pools may include long-lived or inactive molecules that do not necessarily reflect protein turnover or activity, which could be particularly relevant for environments with high levels of reactive oxygen species, such as mitochondria ^40^. The mRNA-protein uncoupling highlights the need for time-resolved measurements of mitochondrial activity rather than relying on steady-state protein abundance alone.

The spatiotemporal aspects of hepatic mitochondrial physiology may also have therapeutic implications, as hepatic AAV-mediated gene addition in mitochondrial disease, or chemically modified RNA interference targeting, will encounter a spatially and temporally heterogeneous transcriptional environment in which target gene expression, and hence therapeutic effect, may differ substantially between PC and PP hepatocytes and feeding-fasting cycles. Incorporating this framework into therapeutic design and preclinical modelling could improve the translational accuracy of such approaches.

In summary, we present a comprehensive dataset describing hepatic mitochondrial regulation that can facilitate analysis, quality control, and annotation of large hepatic datasets, while also providing a foundation for mechanistic and disease-focused studies.

## Data availability

The scRNA-seq data used in this study are available in the Gene Expression Omnibus (GEO) under accession code GSE244392. The GTEx V8 human RNA-seq data used in this study are available in the GTEx Portal database under accession code phs000424.v8.p2, protected data can be requested from the database of Genotypes and Phenotypes (dbGaP). The mouse RNA-seq data generated by Weger et al. and reused in this study are available in GEO with the accession code GSE77221. The human spatial transcriptomics data generated by Yakubovsky et al. and reused in this study are available in Zenodo at https://zenodo.org/records/17735506. The code used for scRNA-seq analysis is available at https://github.com/naef-lab/LiverZonation or from the corresponding author upon reasonable request.

## Acknowledgements

We thank Prof. Felix Naef (EPFL), Prof. Shalev Itzkovitz (Weizmann Institute of Science), Prof. Henrik Oster (University of Lubeck), and Prof. Leonardo Vinícius Monteiro de Assis (University of Gothenburg) for insightful feedback and suggestions on the manuscript. We thank Prof. Graham Knott and the Biological Electron Microscopy Facility (BioEM) at EPFL for scanning electron microscopy. The work was supported by the Slovenian Research Agency (ARIS), grants J3-60069 to T. M. and N1-0305 to U. M. Internal funding from the University of Maribor awarded to T. M. for postdoctoral mentoring of highly prospective researchers covered the salary and work of J. V.; this activity was financially supported under the ARIS stable financing programme SN-ZRD/22-27/0552. The schematic in Fig. 3A was generated with biorender.com.

## Methods

### Analysis of mouse scRNA-seq data

To distinguish *mt-Co1*^+^ and *mt-Co1*^-^cells, *mt-Co1* expression values were extracted from the hepatocyte cluster of the scRNA-seq dataset ^11^ using the *Seurat v5* R package ^41,42^. Cells were classified as positive or negative based on a two-component Gaussian mixture model fitted using the *mclust* R package ^43^, with cells assigned to the high- or low-expressing component, respectively. To better illustrate the clustering of *mt-Co1*^-^cells in SFig. 1B, the cells for genes with higher expression (*Malat1*, *Jun*, *Mtss1*) were not ordered, whereas the cells with lowly expressed genes were ordered, with the cells with highest gene expression depicted at the top layer.

The spatial reconstruction of hepatocytes was previously described in detail ^11^, and the code was made publicly available. The dataset was cross-referenced with MitoCarta 3.0 ^3^ to identify mitochondrial-function associated genes.

### Analysis of mouse bulk RNA-seq data

Preanalysed C57BL/6 bulk RNA-seq data ^27^ were cross-referenced with the mouse MitoCarta 3.0 ^3^ to subset the RNA-seq dataset to mitochondrial-function associated genes. Next, a mitochondrial-function targeted reanalysed was performed with the previously published rhythmicity analysis pipeline, using the *dryR* package ^27^.

### Analysis of mouse proteomics

Time-resolved mouse liver proteomic data, with tissue sampled every 3 h, were obtained from ref. ^2^. Mitochondrial proteins, as annotated in mouse MitoCarta 3.0 ^3^, were retained if they had at most 5 missing values across the samples, and assessed for rhythmicity with the *f_24* harmonic-regression model of the *dryR* package ^27^. A protein was classified as circadian when its rhythmicity p-value was below 0.05 and its relative amplitude was at least 0.2; all other proteins were classified as non-rhythmic. The rhythmicity was visualised with heatmaps, which were generated using *pheatmap* R package from log_2_-transformed mitochondrial protein abundance values across ZT. For visualization, values for each protein were mean-centered across time and row-scaled, such that colors indicate standardised relative abundance within each protein. Consequently, heatmap colors reflect temporal abundance patterns within proteins and are not directly comparable as absolute abundance differences between different proteins.

### Analysis of spatial transcriptomics data

Spatial transcriptomics data from ref. ^12^ (10x Genomics Visium HD, samples M1 and M2) were processed using the *Seurat v5* R library ^41,42^. Filtered count matrices were imported from the 10x HDF5 output with Read10X_h5, spatial coordinates were read from the tissue positions file, and a Seurat object was created with the Spatial assay using the coordinate metadata, while bins outside the tissue were excluded (in_tissue == 1). Mitochondrial content was computed as the percentage of reads mapping to mitochondrial genes. Data were log-normalized to enable downstream feature visualization. For spatial plotting in the native 10x image coordinate system, a custom dimensional reduction was constructed from full-resolution pixel coordinates (*pxl_col_in_fullres*, *pxl_row_in_fullres*) and added to the object as a spatial_map, with the y-axis reversed to match histology image orientation. A region of interest (ROI) was defined by manual lasso selection using Seurat’s CellSelector on the spatial map, selection coordinates were saved for reproducibility, and the dataset was subset to selected bins. The manual selection was used to capture regions with proximal central and portal veins, and a central ROI was chosen to avoid edge effects. Histology overlay was generated by loading the tissue image and scale factors, transforming spot coordinates to image space, and plotting binned locations on top of the H&E raster with *ggplot2*. *Load10X_Spatial* was not used; the spatial image was attached manually by constructing a *Seurat VisiumV1* image object with the H&E raster, scale factors, and coordinate fields formatted to *Seurat*’s expected internal names (imagerow, imagecol), and then stored in the *Seurat* object. Finally, spatial expression patterns for *CYP2E1* and mitochondrial percentage were visualised on the histology background. For display, zero values were set to NA and values were restricted to the 5th–95th percentile range prior to plotting with a *viridis* color scale. For the M1 sample, native 8 µm resolution coordinates were used, whereas for the M2 sample, the 8 µm count matrix was used, and 8 µm bin coordinates were derived from the 2 µm tissue positions by grouping 4 × 4 blocks of 2 µm bins and taking their mean pixel position. To plot spatial expression profiles of individual genes, we used the Human Liver Zonation web applet provided by Yakubovsky et al. (https://itzkovitzwebapps.weizmann.ac.il/webapps) ^12,34^.

### Determination of rhythmicity in human bulk RNA-seq

The GTEx v8 liver bulk RNA-seq dataset was restricted to human MitoCarta 3.0 mitochondrial genes ^3^, yielding 1007 genes across 224 donors (160 men, 64 women). Rhythmicity in the dataset was assessed using a previously developed pipeline ^37^. Briefly, counts were TMM-normalised to log_2_ CPM with *edgeR* ^44^, and per-gene variance from sex, age, ischaemic time and Hardy death-class was regressed out. Each donor’s circadian phase (donor internal phase, DIP) was taken unchanged from ^37^, where *CHIRAL* infers one phase per donor from twelve clock transcripts across all tissues, anchored to skeletal muscle (https://github.com/naef-lab/CHIRAL) ^37^. According to the pipeline, per sex, a 24-h cosinor (harmonic) model was fitted against the DIP, with significance by likelihood-ratio test against an intercept-only model. Peak-to-trough amplitude and phase were derived from the fitted cosine. Differential rhythmicity between male and female donors was assessed as previously described ^27,37^. A gene was assigned to model 0 (not robustly assessable) if it failed to meet both q(BH) < 0.2 and a log_2_ peak-to-trough amplitude > 0.5 in all of the male, female and pooled (sex-combined) harmonic fits; otherwise it was classified as model 1, non-rhythmic in both sexes; model 2, rhythmic in men only; model 3, rhythmic in women only; model 4, rhythmic in both sexes with shared phase and amplitude; or model 5, rhythmic in both sexes with sex-dimorphic phase and/or amplitude.

### Electron microscopy

TEM and SEM experiments were previously performed and the method described in detail ^11^. The tissue was collected at ZT 2. Here, the data were reanalysed for 2D and 3D visualisations. For 3D analysis, brightness and contrast normalisation between planes was performed to reduce slice-to-slice variability before segmentation. Two complementary approaches were used. In the first approach, the mean gray level of each slice was computed. For adjacent slices, pairwise mean gray levels were calculated, and slice pairs with ratios below 98 percent or above 102 percent were identified as intensity outliers. For outlier slices, their gray levels were corrected by setting the mean outlier gray level to the ratio of 1, compared to the reference/previous slice (RStudio 2025.09.1 build 401, R version 4.5.1). In the second approach, slice intensities were normalised using histogram matching (Fiji, v 1.54p). This method adjusts the full gray level distribution of each slice to match a reference slice, which improves interslice consistency in a process based on the histogram harmonisation strategies ^45^. Together, these steps reduced brightness and contrast fluctuations that might otherwise affect region delineation or quantitative analysis.

For SEM reconstruction, a 2.5D U-Net deep-learning segmentation model (depth level 5, initial filter count 65, input slice count 3) was applied, and trained from scratch using the Segmentation Wizard in Dragonfly 3D World (Version 2024.1 build 1601; Comet Technologies Canada Inc, Canada). Five classes were defined: mitochondria, lipid droplets, nucleus, cell area, and all remaining unlabelled voxels. The initial training dataset consisted of 10 manually segmented frames, while an additional dataset was trained using 5 pre-segmented frames. Training was performed for 100 epochs with data augmentation set to 10, using the Categorical Cross-Entropy loss function and the Adadelta optimisation algorithm. The input voxel size was 7 × 7 × 50 nm^3^. To reduce the risk of false positive detections and the likelihood of noise-driven microregions, all connected components smaller than 10 voxels were removed ^46^.

---

**Supplementary Figure 1:**
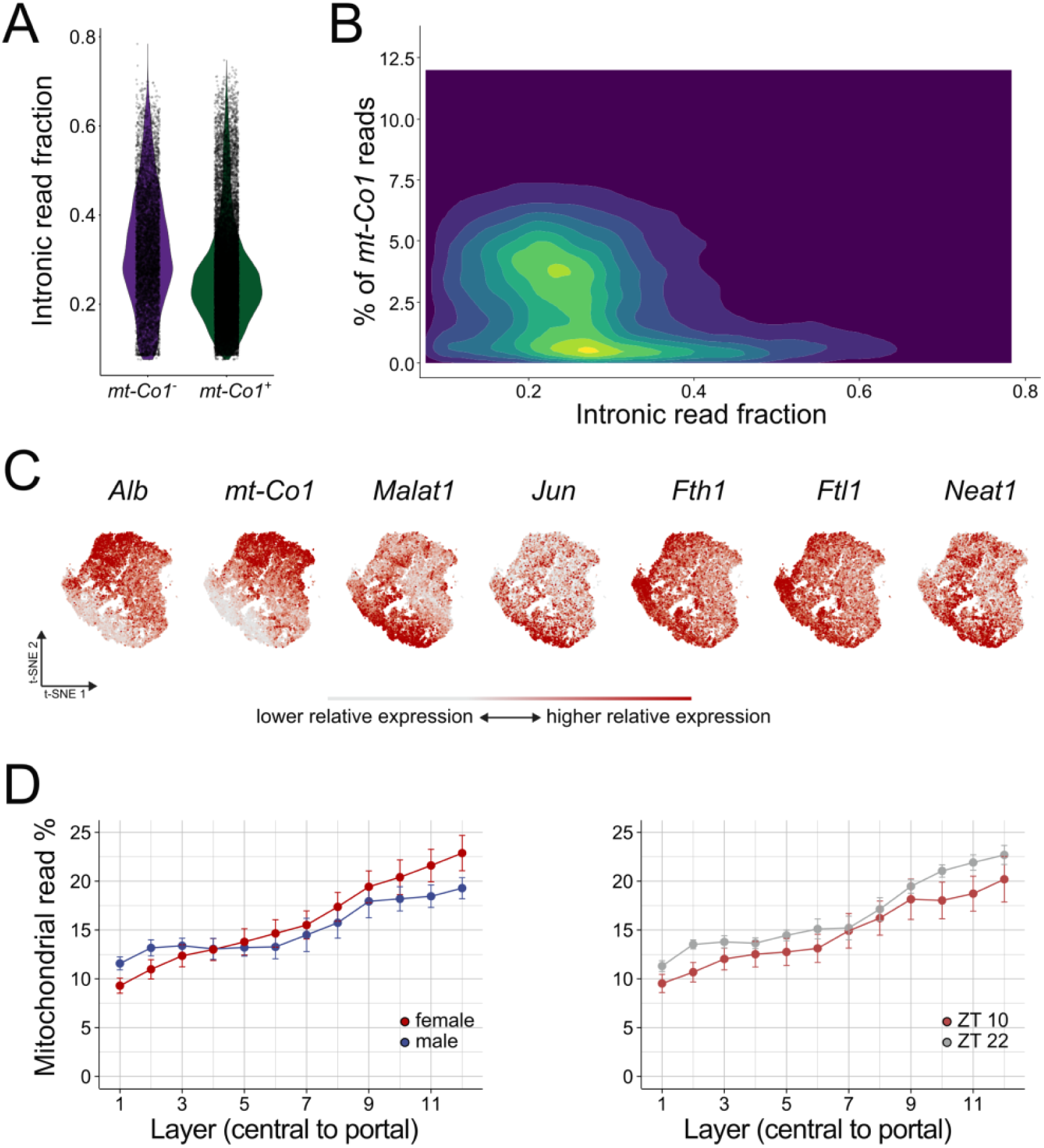
Characteristics of mt-Co1^⁻^ cells in 129S mice. (A) Both mt-Co1^⁻^ and mt-Co1^+^ cells show a comparable intronic read fraction. (B) Two-dimensional density plot of the percentage of mt-Co1 reads per cell versus intronic read fraction. Two high-density populations are apparent, corresponding to cells with low and high mt-Co1 read fractions. (C) Dimensionality reduction of hepatocytes shows a higher expression of Malat1, Jun, Fth1, Ftl1 and Neat1 in the hepatocytes that show low mt-Co1 expression. (D) Aggregated visualisation of mitochondrial read fraction across the porto–central axis, split by sex (left panel) and time of tissue isolation (right panel), based on data from ref. ^11^.

**Supplementary Figure 2:**
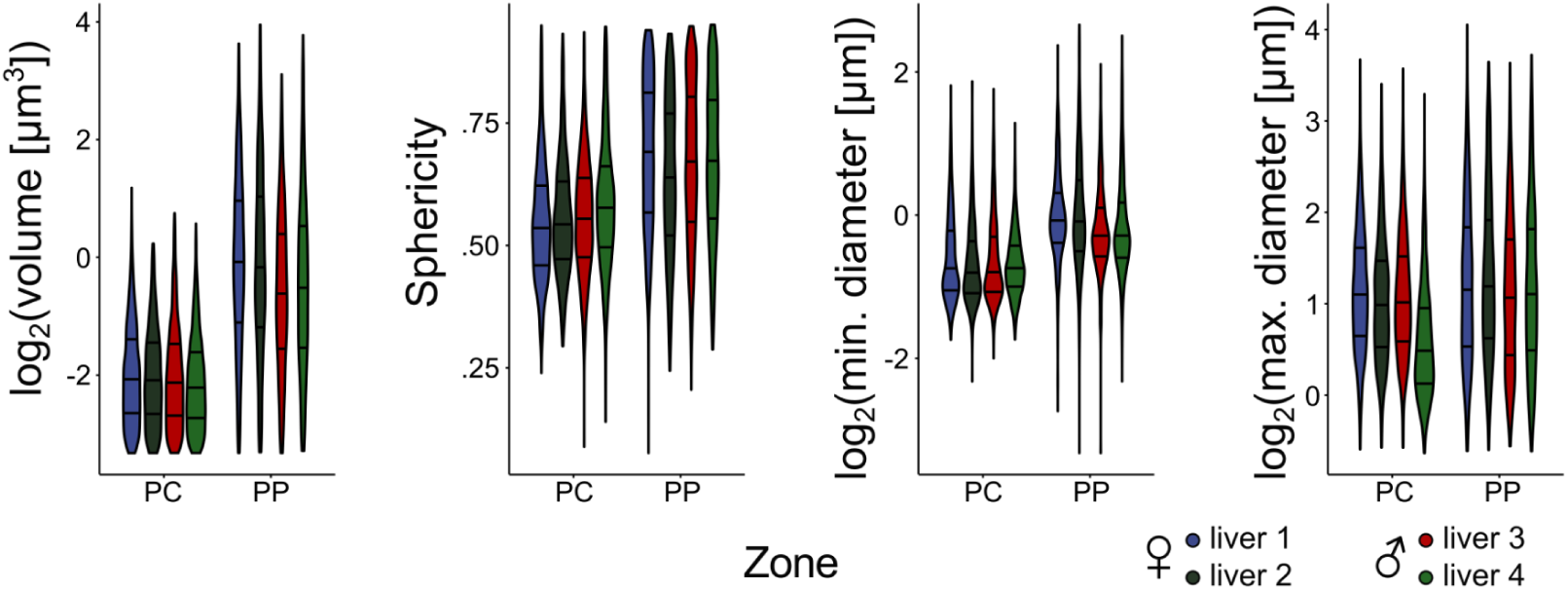
Morphological characteristics of pericentral and periportal hepatocytes in 129S mice. The measured morphological characteristics are presented on a per-sample basis, where the violin plot shows the distribution of measurements of individual mitochondria. The results demonstrate reproducibility between biological replicates. The volume, sphericity and min. and max. Feret diameters are shown for each sample.

**Supplementary Figure 3:**
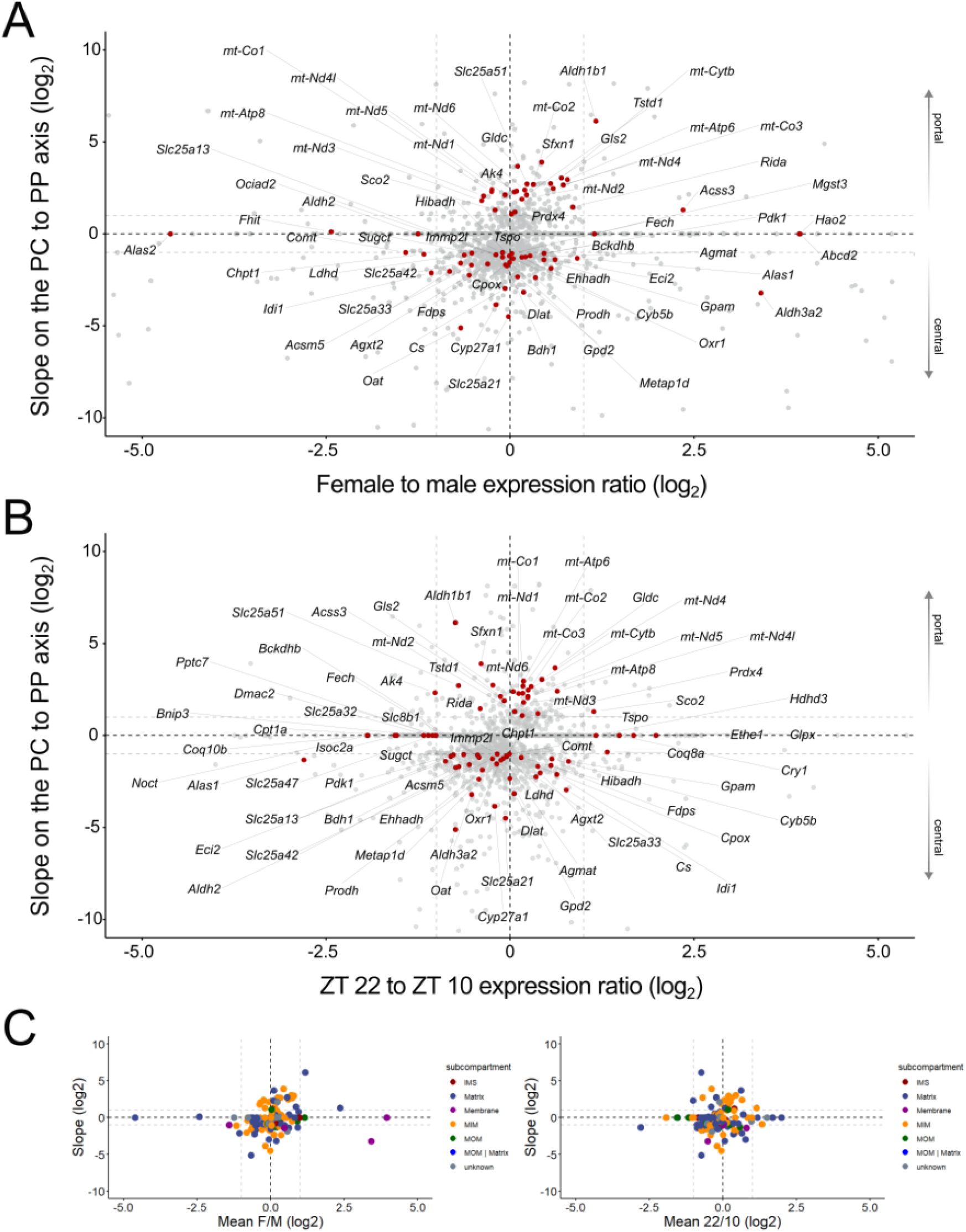
Expression of mitochondrial genes in spatially reconstructed single-cell data in 129S mice. (A-B) Zonation is represented by the average slope (negative slopes characterise central profiles) across all conditions (y-axis) vs. (A) the female to male expression ratio or (B) ZT 22 to ZT 10 expression (bottom panel). Genes in the corners are highly zonated and sexually dimorphic or highly zonated and strongly regulated by time. All mitochondrial genes (as annotated in MitoCarta 3.0), whose log2FC changes > 1 with zone, sex or time are marked with red dots and explicitly named. The plots show that key mitochondrial metabolic enzymes are not only spatiotemporally regulated, but also sexually dimorphic, demonstrating complexity of mitochondrial function. E.g., the very long chain fatty acyl CoA transporter Abcd2, the mitochondrial short chain acyl CoA synthetase Acss3, and Fech, which catalyzes the last step of heme biosynthesis, show markedly increased expression in female livers. Conversely, Idi1, a critical enzyme in the mevalonate/cholesterol biosynthesis pathway, exhibits markedly higher male expression. (C) Analogous plots to those shown in panels A and B, with mitochondrial genes labelled by mitochondrial subcompartment, showing that no clear subcompartment-wide bias was apparent.

**Supplementary Figure 4:**
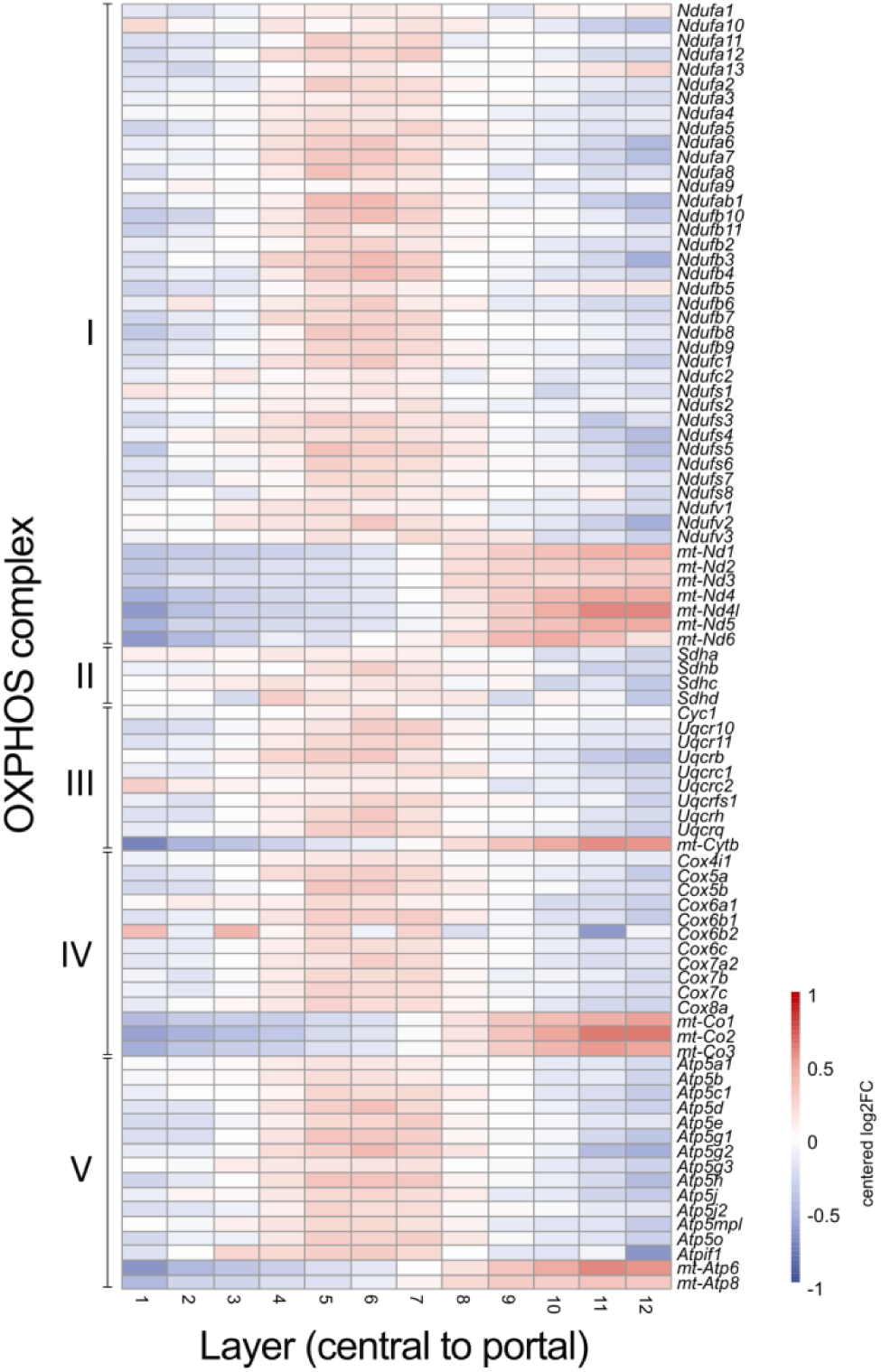
Zonated expression of OXPHOS genes in 129S mice. Zonation is represented by 12 virtual layers of hepatocytes, where 1 represents the most PC layer and 12 the most PP layer. Per-gene expression is depicted as row-centered log_2_-fold changes, i.e. for each layer the log_2_-fold deviation from the mean expression across all layers, capped at –1 and 1, respectively.

**Supplementary Figure 5:**
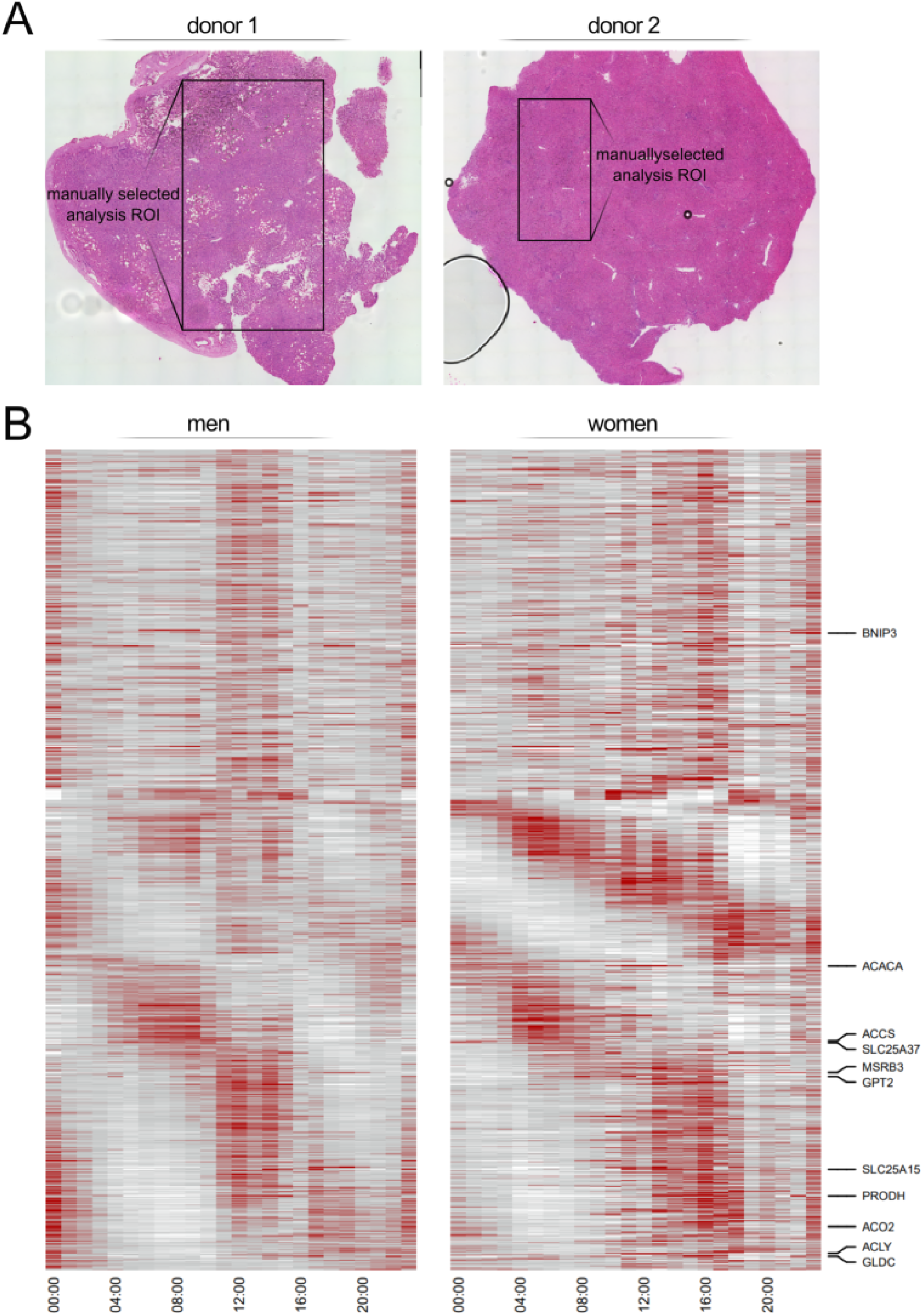
(A) Visium HD sample overview. The full histologically stained tissue overview from ref. ^12^ is shown, with the box depicting the region of interest (ROI), which we manually selected and used for the analysis. The manual selection was used to capture regions with central and portal veins, and a central ROI was chosen to avoid edge effects. The cells on sample edges experience damage and cytoplasm leakage, artificially increasing the mitochondrial read fraction. (B) Analysis of mRNA expression for mitochondrial genes in human bulk RNA-seq data from the GTEx project. The heatmaps show genes that could not be statistically assessed for rhythmicity in men (left panel) and women (right panel; n = 224; 160 men, 64 women).

