## Supplementary Figures for "The hepatic mitochondrial landscape"

### Supplementary Figures: The hepatic mitochondrial landscape

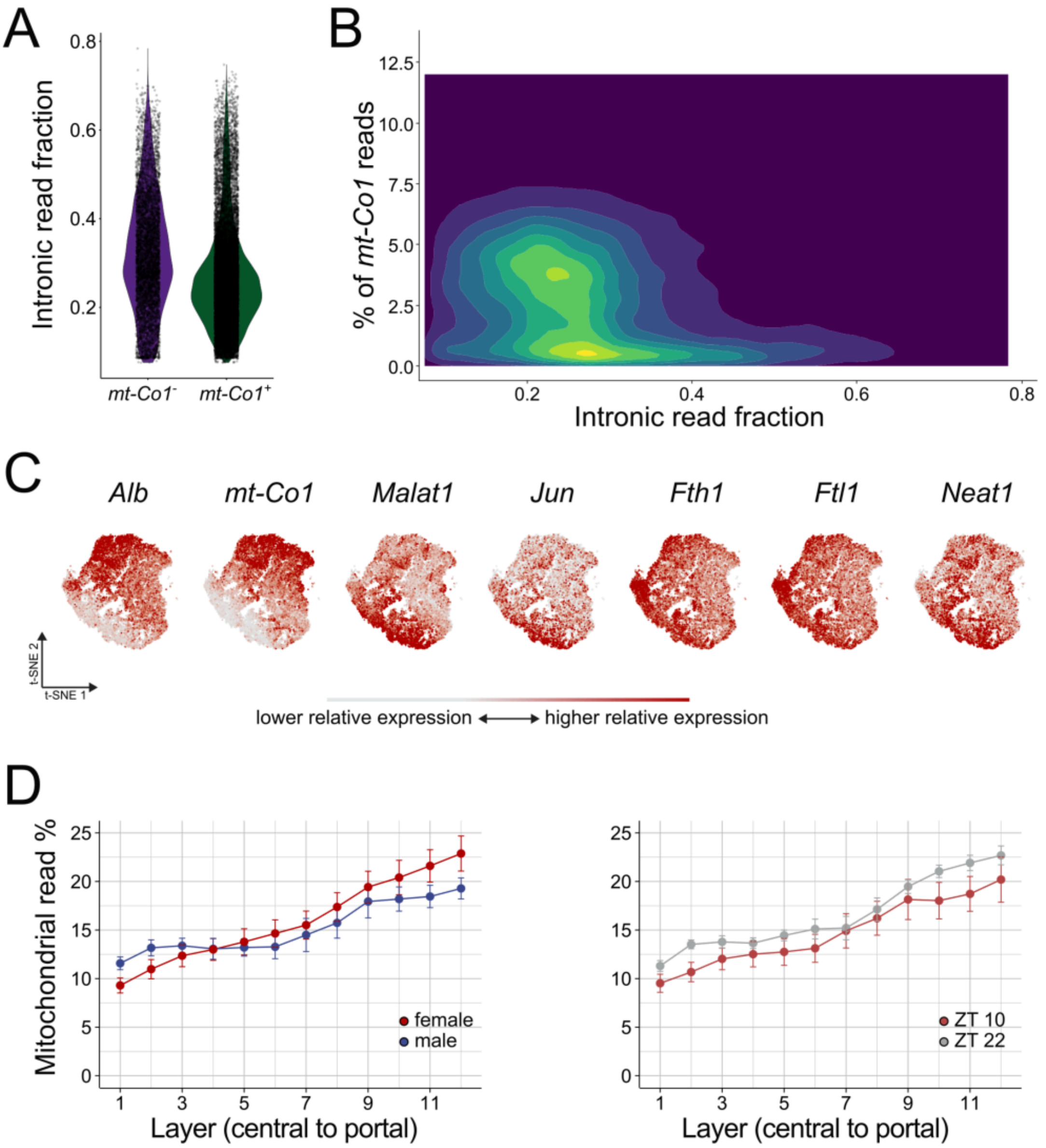


*Supplementary Figure 1:* ***Characteristics of mt-Co1$^-$ cells in 129S mice.*** *(A) Both mt-Co1$^-$ and mt-Co1$^+$ cells show a comparable intronic read fraction. (B) Two-dimensional density plot of the percentage of mt-Co1 reads per cell versus intronic read fraction. Two high-density populations are apparent, corresponding to cells with low and high mt-Co1 read fractions. (C) Dimensionality reduction of hepatocytes shows a higher expression of Malat1, Jun, Fth1, Ftl1 and Neat1 in the hepatocytes that show low mt-Co1 expression. (D) Aggregated visualisation of mitochondrial read fraction across the porto–central axis, split by sex (left panel) and time of tissue isolation (right panel), based on data from ref.* [10].

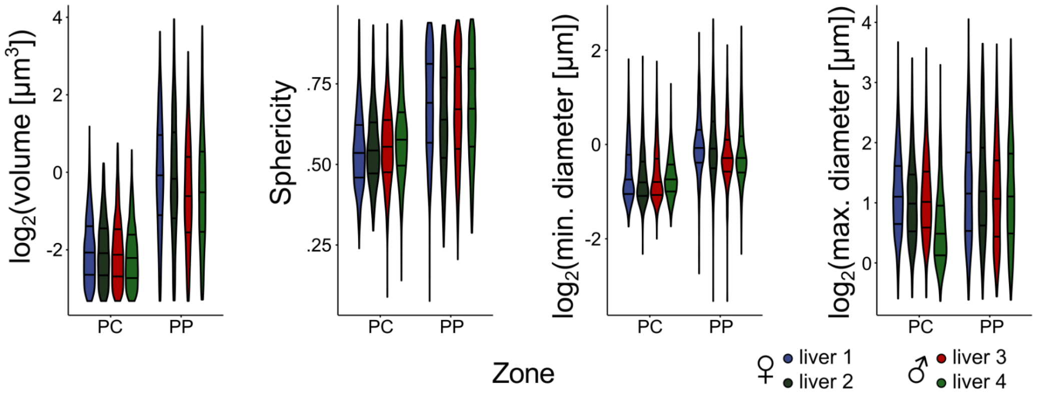


*Supplementary Figure 2:* ***Morphological characteristics of pericentral and periportal hepatocytes in 129S mice.*** *The measured morphological characteristics are presented on a per-sample basis, where the violin plot shows the distribution of measurements of individual mitochondria. The results demonstrate reproducibility between biological replicates. The volume, sphericity and min. and max. Feret diameters are shown for each sample.*

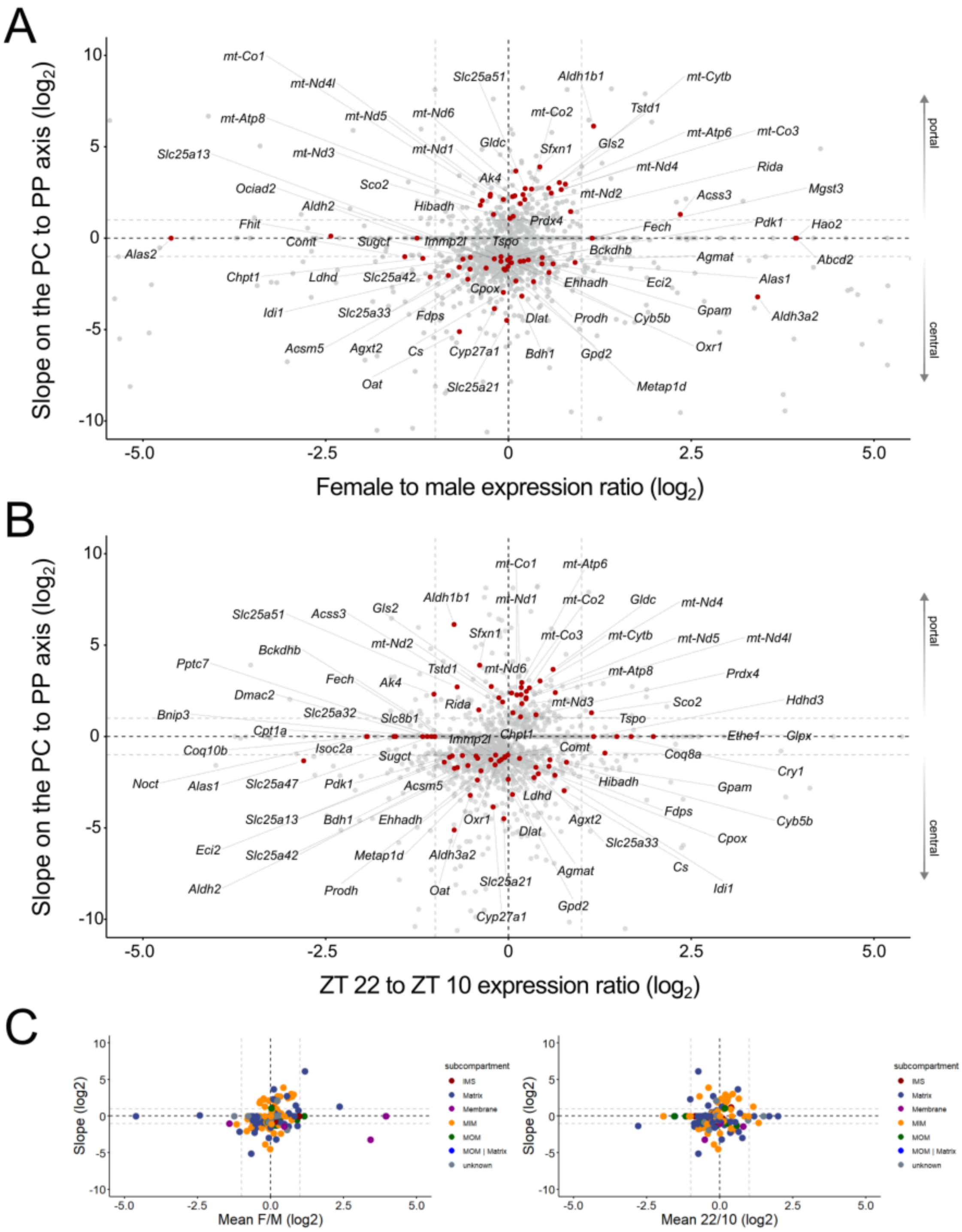


*Supplementary Figure 3:* ***Expression of mitochondrial genes in spatially reconstructed single-cell data in 129S mice.*** *(A-B) Zonation is represented by the average slope (negative slopes characterise central profiles) across all conditions (y-axis) vs. (A) the female to male expression ratio or (B) ZT 22 to ZT 10 expression (bottom panel). Genes in the corners are highly zonated and sexually dimorphic or highly zonated and strongly regulated by time. All mitochondrial genes (as annotated in MitoCarta 3.0), whose log2FC changes > 1 with zone, sex or time are marked with red dots and explicitly named. The plots show that key mitochondrial metabolic enzymes are not only spatiotemporally regulated, but also sexually dimorphic, demonstrating complexity of mitochondrial function. E.g., the very long chain fatty acyl CoA transporter Abcd2, the mitochondrial short chain acyl*

*CoA synthetase Acss3, and Fech, which catalyzes the last step of heme biosynthesis, show markedly increased expression in female livers. Conversely, Idi1, a critical enzyme in the mevalonate/cholesterol biosynthesis pathway, exhibits markedly higher male expression. (C) Analogous plots to those shown in panels A and B, with mitochondrial genes labelled by mitochondrial subcompartment, showing that no clear subcompartment-wide bias was apparent.*

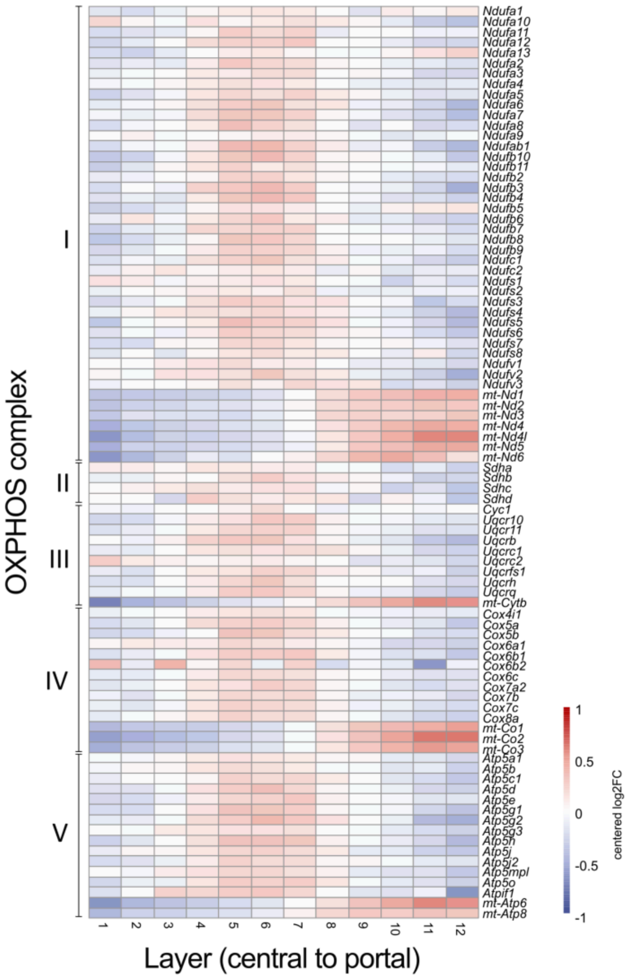


*Supplementary Figure 4:* ***Zonated expression of OXPHOS genes in 129S mice.*** *Zonation is represented by 12 virtual layers of hepatocytes, where 1 represents the most PC layer and 12 the most PP layer. Per-gene expression is depicted as row-centered log*$_2$*-fold changes, i.e. for each layer the log*$_2$*-fold deviation from the mean expression across all layers, capped at -1 and 1, respectively.*

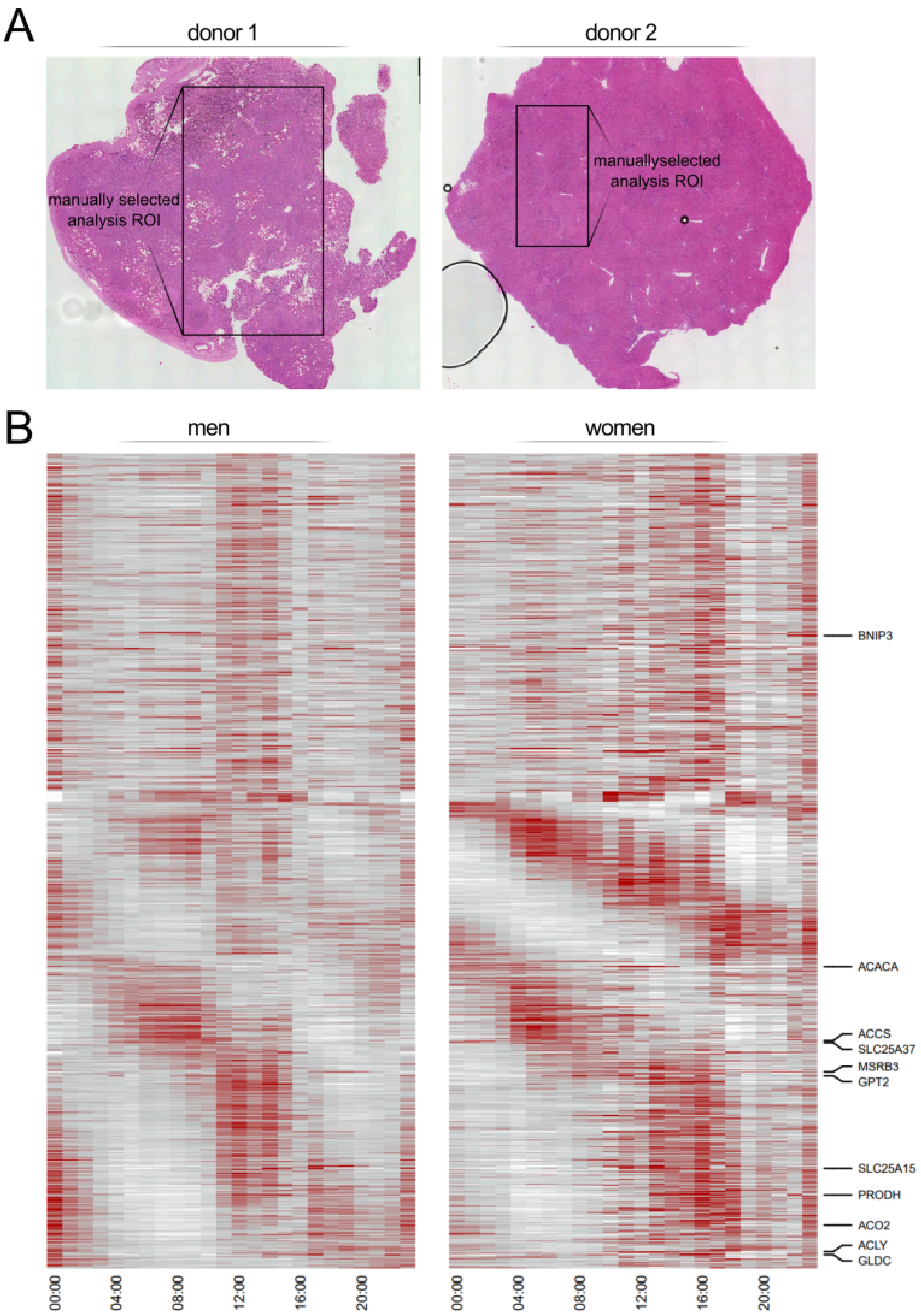


*Supplementary Figure 5: (A) Visium HD sample overview. The full histologically stained tissue overview from ref. [19] is shown, with the box depicting the region of interest (ROI), which we manually selected and used for the analysis. The manual selection was used to capture regions with central and portal veins, and a central ROI was chosen to avoid edge effects. The cells on sample edges*

*experience damage and cytoplasm leakage, artificially increasing the mitochondrial read fraction. (B) Analysis of mRNA expression for mitochondrial genes in human bulk RNA-seq data from the GTEx project. The heatmaps show genes that could not be statistically assessed for rhythmicity in men (left panel) and women (right panel; n = 224; 160 men, 64 women).*
